# GMSECT: <u>G</u>enome-Wide <u>M</u>assive <u>S</u>equence <u>E</u>xhaustive <u>C</u>omparison <u>T</u>ool for structural and copy number variations

**DOI:** 10.1101/2021.03.01.433223

**Authors:** Abhishek Narain Singh

**Affiliations:** University of Eastern Finland

**Author notes:** Web: ABioTek www.tinyurl.com/abinarain.

## Abstract

GMSECT is a parallel robust ‘*Application Interface*’ that efficiently handles the large genomic sequences for rapid and efficient processing. It is a ‘message passing interface’ based parallel computing ‘Tool’ that can be operated on a cluster for ‘Massive Sequences Exhaustive Comparison’, to identify matches such as the structural variants. The GMSECT algorithm can be implemented using other parallel application programming interfaces as well such as Posix-Threads or can even be implemented in a serial submission fashion. There is complete flexibility to the choice of comparison tool that can be deployed and with the optional parameters as of the choice of comparison tools to suit the speed, sensitivity and specificity of pair wise alignment. The algorithm is simple and robust, and can be applied to compare multiple genomes, chromosomes or large sequences, of different individuals for personalized genome comparison and works good for homologous as well as distant species. The tool can even be applied to smaller genomes like the microbial genome such as the *Escherichia coli* or algae such as *Chlamydomonas reihardtti* or yeast *Saccharomyces cerevisiae* to quickly conduct comparisons, and thus finds its application to the pharmaceuticals and microbial product based firms for research and development. The application Interface can efficiently and rapidly compare massive sequences to detect for the presence of numerous types of DNA variation existing in the genome ranging from single nucleotide polymorphism (SNPs) to larger structural alterations, such as copy-number variants (CNVs) and inversions. The new algorithm has been tested for comparing the chromosome 21 of Celera’s R27c compilation with all the 48 chromosomes of Celera’s R27c compilation and with all the 48 chromosomes of the human Build 35 reference sequence, which took just 2 Hours and 10 minutes using the pair wise blast algorithm choice and with 110 processors each with 2.2 GHz capacity and 2 GB memory. GMSECT facilitates rapid scanning and interpretation in personalized sequencing project. The application interface with the above resources and alignment choice is estimated to do exhaustive comparison of the human genome with itself in just 2.35 days. An exhaustive comparison of an individual’s genome with a reference genome would comprise of a two ‘self genome’ comparison and a ‘non-self genome’ comparison which is estimated to take about 9.4 days with the above resources. With the advent of personalized genome sequencing project, it would be desirable to compare 100s of individual’s genome with a reference genome. This would involve a ‘non-self genome’ and a ‘self genome’ comparison for each genome, and would take around 7 days for each individual’s genome using GMSECT and the above mentioned resources.

## Introduction

Sequence homology has been an established approach to get the first hint of functional similarity. As the DNA sequence database continues to grow, the chances of getting closer matches keeps on improving. However, with the increase in the DNA repository, there is an equal increase in demand for handling the sequences in an efficient and rapid way. Most of the existing pair wise alignment tools are an extension to the dynamic programming algorithm, and though they are extensively fast in comparison to standard dynamic programming approach, they are not rapid and efficient to handle massive sequences, resulting in memory address violation and other computational complications. GMSECT takes into consideration of memory and processor resources to efficiently do the alignment comparison with minimal time using parallel computing approach. GMSECT is an application interface which can be used with any pairwise alignment tool such as Blast, Blat, Fasta or any other possible alignment tool.

### What are the massive sequences?

Massiveness of a sequence is dictated not just in terms of the size of the genome, but the computational complexity that increases with incremental size. In other words, a massive sequence is one that takes a lot of time to process using standard computational resources, and many a times do not get processed reasonably well. Massive sequences are the sequences which are characteristics of higher eukaryotes such as the vertebrates and the plants which have their genome size in the order of hundreds of megabases. Typical example includes the model plant *Arabidopsis thaliana* genome and the Human genome. For instance, the size in megabytes of chromosome 1, 2 and 3 of human Build 35 reference sequence are 241MB, 237MB and 195MB respectively. However, the term massive is relative and should be used in context with the algorithm implemented. One reason for the prior statement is the difference in the number of words generated by different algorithms and thereby the hits resulting in high scoring pairs (HSPs) and their extension until the score drops below the set threshold value. Thus there is a computational limit with respect to the memory resource. The compared sequences should not be so massive such as to cause ‘memory address violation’, resulting in ‘core’ files generation or other errors such as ‘segmentation fault’, ‘mpid:Broken pipe!’, or a cause of the machine to ‘hang’. For instance, while the massiveness limit on a 2GB, 2.2GHz Processor for *blastn* could be around 50,000 bases, *blat* 20,000 bases. Of course each of these heuristics has their own merits and demerits under various requirements to have different suitability. Under fair approximation, *‘number of bases could be considered roughly to be equal to number of bytes’*. Because of the massive size of sequences and computational requirements, the different pair wise alignment algorithms are impractical without the use of a supercomputer. The present algorithm serves as a parallel computing interface to the existing heuristic tools which can be operated on a cluster of processors.

### Speed, sensitivity and specificity

The variants of dynamic programming algorithm fall under the umbrella of *Needleman Wunsh* Algorithm for global alignment and *Smith-Waterman* algorithm for local alignment. A popular local alignment tool such as blast, works by looking for matches of all possible words of size ***w*** and match score threshold ***T*** or more, and then extending the matches by dynamic programming until the score drops below the threshold value ***T***. The number of words generated for nucleotides is 4^***w*** since there are four bases viz., T, C, A & G. While a small value of ***w*** would generate fewer words but result in high number of HSPs (High Scoring Pairs), high value of ***w*** would generate more words but result in less HSP. Hence in the former case there is high sensitivity, while the latter case provides more specificity. With an increase in sensitivity there are problems of high noise and thus redundant information, whereas with an increase in specificity there are concerns for missing out relevant matches. For instance, as an extreme case example if ***w=1*** we generate 4^1 i.e., just 4 words viz., T, C, A & G. However, all of these four words would generate HSPs to match up the entire sequences, thereby generating huge amount of redundant data consuming high computational time. As a converse extreme example, say we were to compare a sequence of size ‘N’ with itself and the word size is kept ***w=N***. This would result in a large number of words to be generated 4^N, but only one of these words would be creating a HSP, thereby resulting in loss of all possible intra-sequence matches such as the Copy Numbers, Inversions, LINES, SINES, mini-satellites, micro-satellites and SNPs. The story is the same for protein sequence matches with a slight modification that now the number of words would be 20^w since there are 20 naturally occurring amino acids. Different alignment algorithms are thus suitable for different data quality since they have their ***w*** and ***T*** set to certain value. It is to be noted that most of the time consumption is in the extension of the HSPs. The speed, sensitivity and specificity of an algorithm on a given dataset are a function of **data quality**, ***w*** and ***T***.

Although there is a linear relationship between the number of words generated and execution time, the number of word hits increases exponentially with decreasing ***T*** (Atshul et. al., J.Mol. Biol. (1990)215, 403-410).

### Data Quality

The DNA sequence is ***non-random***. As an example we know of the presence of CpG islands or the locally biased A and T rich region. More GC percentage is known to be associated with more DNA stability, thermal stability, and species evolve with codon biasing due to stability criteria or some other forces. The mutation, addition or deletion of single nucleotides or large chunks of DNA undergo a survival selection test for its existence. The selection check depends on the existing metabolic pathway in a cell, because of which distant species have different selection criteria to any change in the genomic sequence. These phenomena suggests as of why does the data quality of a distant species, say, *Arabidopsis thaliana* is different from human beings. The survival criteria and complexity in higher organism is different than the lower organism, because of the existing metabolic pathways, and thus while a prokaryote’s ORF does not have introns, the eukaryotic genomic sequence is segmented by splice sites as introns and exons. Further, while the average gene length in prokaryotes is about 1Kbp the genes in eukaryotes can be as big as 15Kbp. We do not yet fully understand the Genome Complexity and thus some researchers consider dusting and masking the ‘uninformative’ region in the genome such as the tandem repeats and fingerprints before making comparisons. Whatever be the case of handling the genomic data, it is for sure that Genome sequence of different organism will have different data quality and thus influence the number of Hits of words generated and thereby the extension as well, resulting in variation in execution time.

### Time Complexity for Aligners

The time computational complexity of pairwise alignment is approximately, ***aW + bN***_***2***_ ***+ c N***_***2***_***W/20^w***, (Atshul et. al., J.Mol. Biol.1990)215, 403-410),

Where, W is the number of words generated, *N*_*2*_ is the number of residues in the subject database and a, b and c are constants. The above formula, would have the number 20 replaced by 4 if the sequence comparison was for nucleotides, and then *N*_*2*_ would be the number of bases in the subject sequence. Also, let *N*_*1*_ be the number of bases in the query. Although the number of words generated, W, increases exponentially with decreasing **T**, it increases only linearly with the length of the query, so that doubling the query length doubles the number of words, (Atshul et. al., J.Mol. Biol. (1990)215, 403-410). That is to say for a given *data quality*, keeping **w** and **T** to be constant, **w= d *N*_*1*_**, where d is proportionality constant.

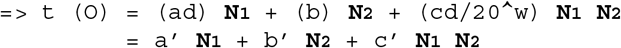

Thus, **t (O) = O (N**_**1**_ **N**_**2**_) i.e., query and subject sequences product.

### Personalized Genome Comparison

As we enter into the era of personalized genome sequencing, we would require a more handy and powerful tool to extract meaningful information from the sequence so as to relate the individual with genetic causes of diseases, such as autism, or in order to understand the genetic cause of a trait of an individual. This would require a reference genome to be set as a standard, to which the individual’s genome could be compared to. We would require a onetime comparison of the reference genome with itself. Additionally, each individual genome would be required to compare with itself as well as with the reference genome. However, the choice of standard reference genome is questionable itself, as no individual’s genome can be biased to be assigned as a reference genome, for the simple reason that a single person is not completely representative of all possible variants in the course of human evolution. The idea should be to create a hypothetical reference genome by extracting statistically relevant information from as many genomes as possible from population of varied races. This statistically relevant information would involve the different structural variants in the genome, such that all major structural variant are being incorporated into the hypothetical genome. Of course, as more and more individual genome would be made available from varied population, the structure of the reference genome would also need to be updated. If the sequencing technology becomes rapid enough, we would be expecting more and more of individual’s genome being available. Ideally, the reference genome should be dynamically updated, though of course with high computational requirement. The dynamic update can be made discrete by making the update periodical and thus reducing computational demand.

For many clinical purposes we choose mouse to be the candidate animal for carrying out experiments on it. Scientists would be interested in knowing the identity of an individual’s genome with the standard mouse genome with respect to the structural variants. Scientists are also interested in the structural variants of a chimpanzee genome to that of human. Likewise, there is a need of comparing two close as well as two distant species in order to understand the genetic makeup, evolution, and to target disease susceptibility.

In fact, the research and development section of many pharmaceutical firms also frequently require finding out the alignment matches of microbes to progress with the experimental bench work. GMSECT can be made use to cut short the time by many a fold.

If we were to compare two genomes, then we would require two ‘*self comparisons*’ of the genomes with itself, and a ‘*non-self comparison*’ of the genomes with each other.

### Non-self comparison

A non self comparison of two genome sequences would involve all versus all chromosome comparison such as filling up the entire matrix. In figure 1 is a schematic comparison.

**Figure 1.**
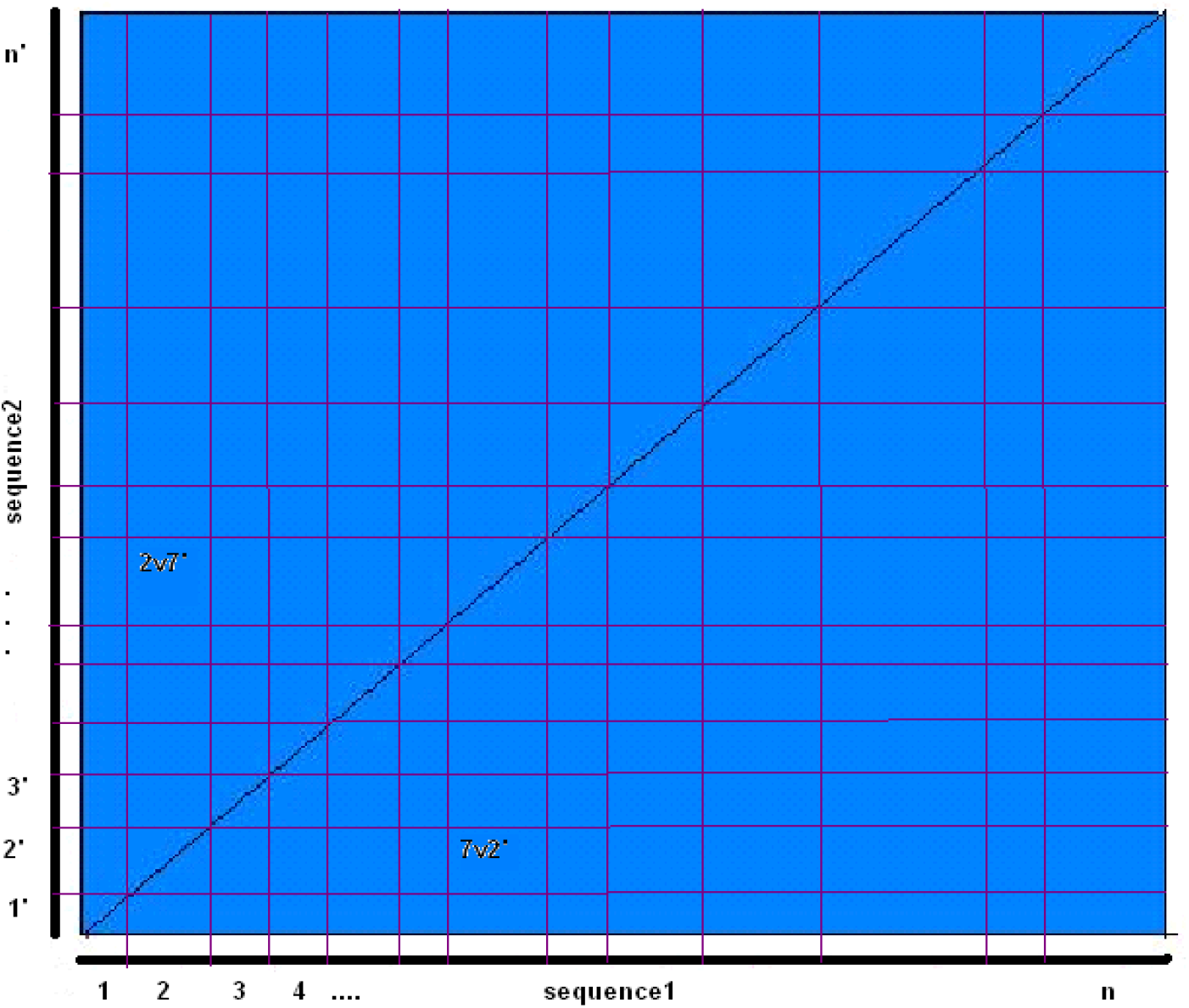

### Self comparison

A self comparison of a genome sequence of an organism or individual with itself would essentially generate a symmetric matrix, such that we would just require the diagonal element comparisons and half of the remaining matrix comparisons. In figure 2 is a schematic view of what would be required.

**Figure 2.**
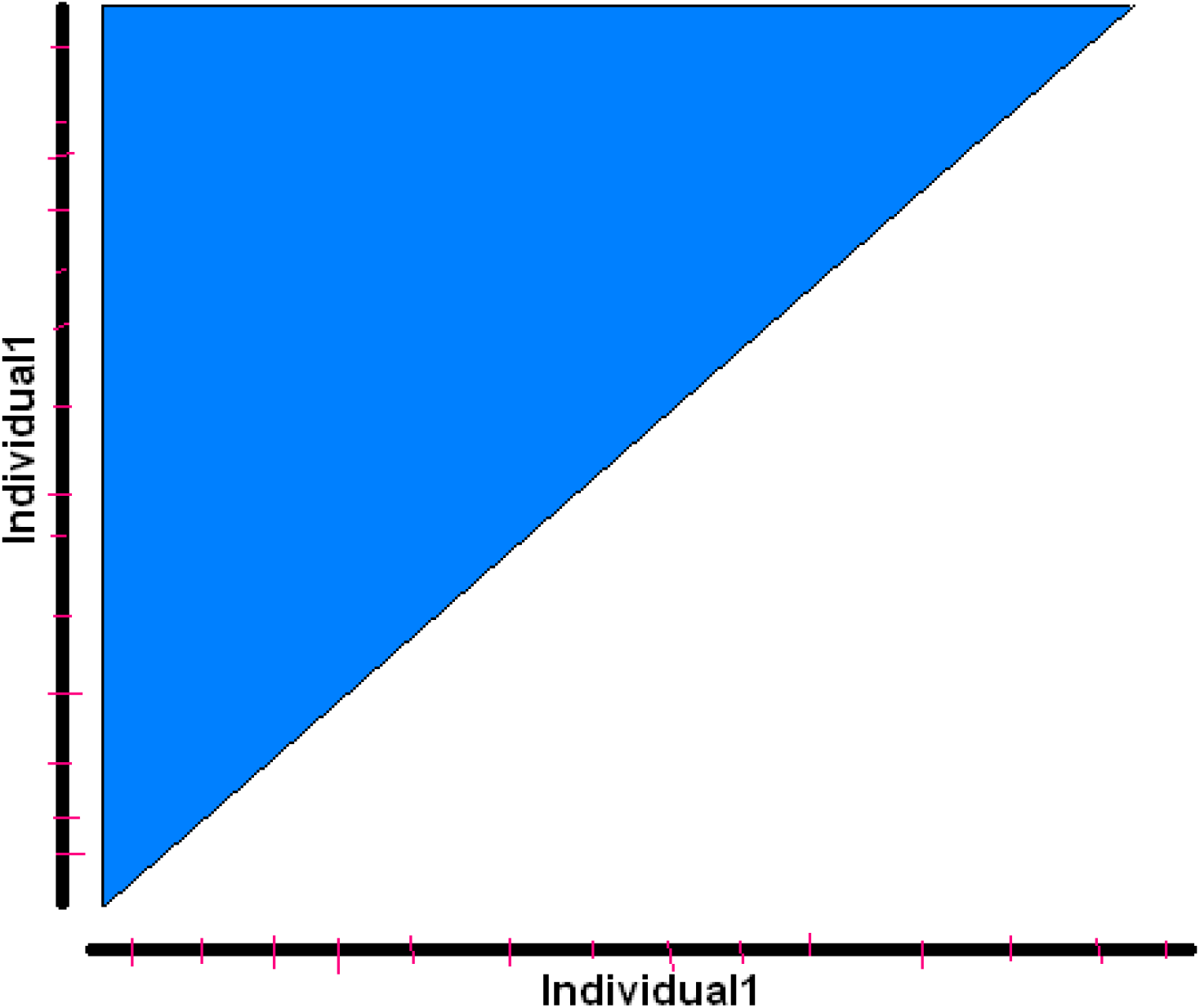

### Genome Alignment Strategy

In order to obtain a ‘non-self comparison’ genome and two ‘self comparison’ of genome the genome should be aligned for comparison in much the same way as we align two sequences. In figure 3 is shown a faulty alignment followed by the right alignment strategy in figure 4.

**Figure 3.**
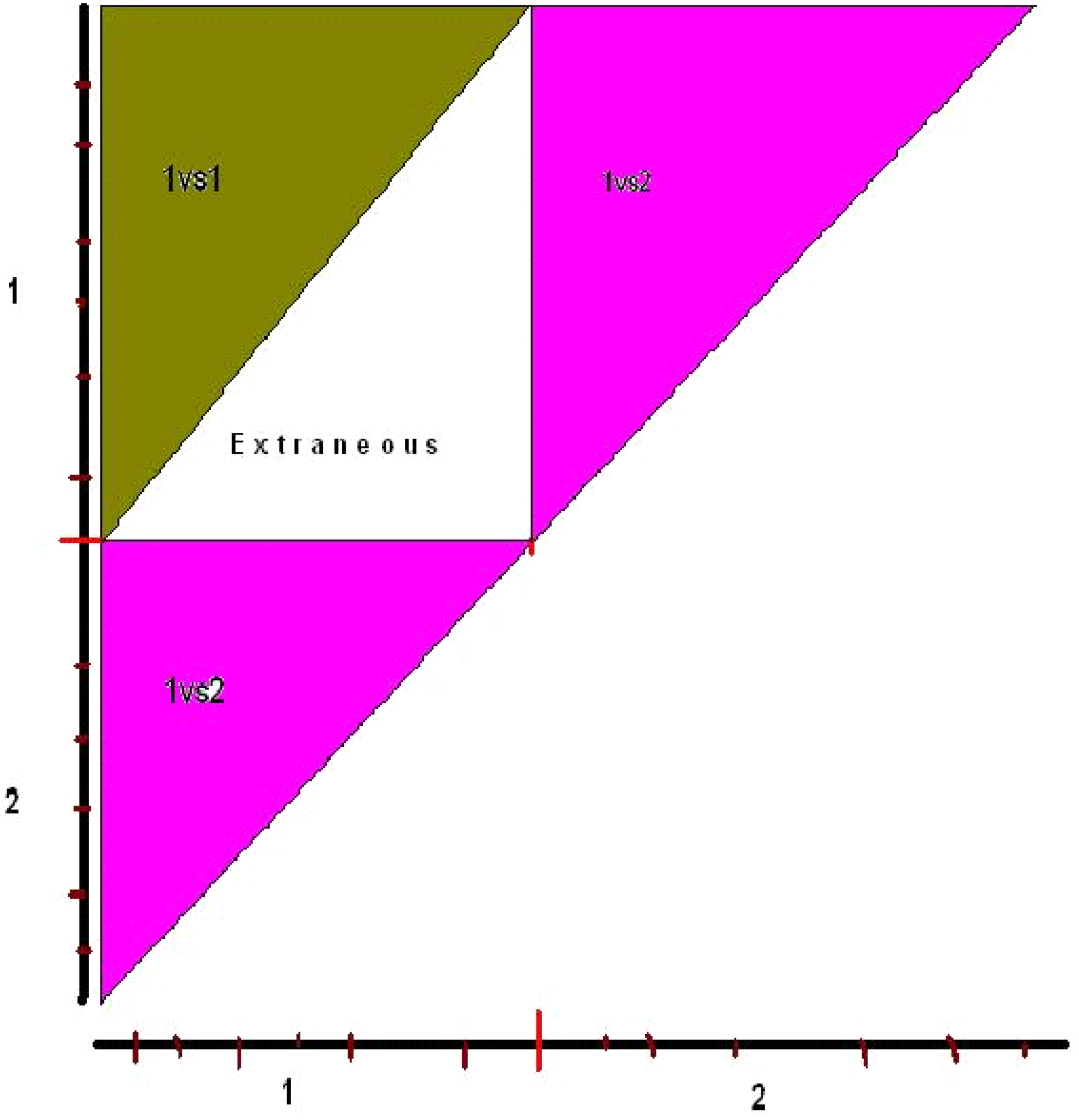

**Figure 4.**
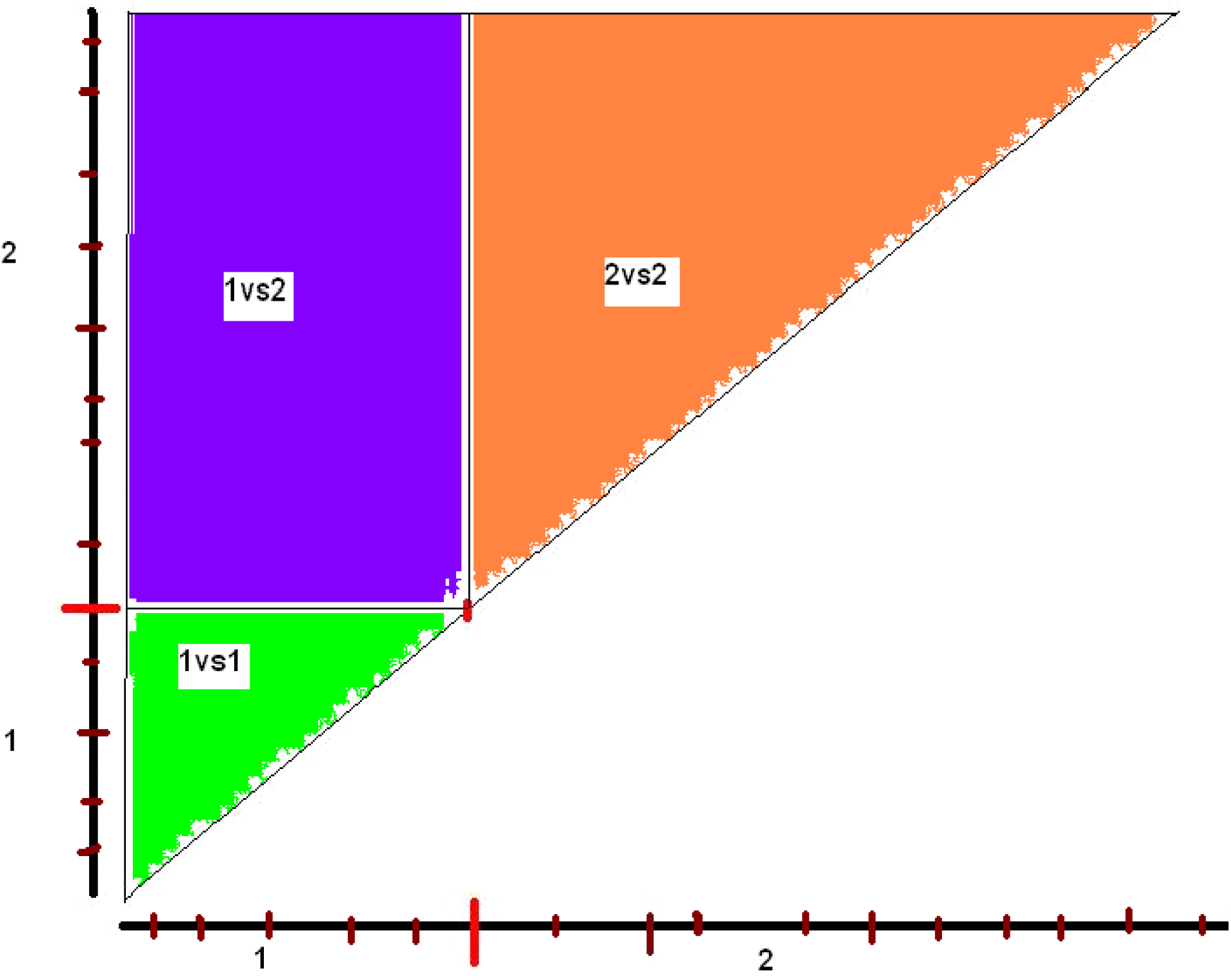

The above faulty alignment misses out the genome2 versus genome2 comparison, apart from doing extraneous computation for genome1 versus genome1. Note that, ‘non-self comparison’ of genome1 versus genome2 is complete, though the square matrix is split up into two triangles at different positions with repetition of the diagonal element comparisons.

The above genome alignment strategy takes care of the ‘self comparison’ as well as the ‘non-self comparison’ for two different genomes. The above strategy works well for comparing genomes of even different sizes, say if one would like to compare the mouse genome with the human genome.

### Divide and Rule (Fragmentation)

For our analysis purpose we took the first individual genome to be the human Build 35 reference genome sequence, such that its 24 chromosomes are numbered as chrP1 to chrPY. We took the second individual genome to be the Celera’s R27c compilation of human genome, such that its 24 chromosomes are numbered as chrC1 to chrCY. A ‘non-self comparison’ of the genomes would represent the graph of the square matrix discussed above.

However, as we pointed out earlier, the chromosomal sequences are massive, and their sizes are in the order of hundreds of megabases. Different comparison heuristic algorithms have different massive sequence limits that they can operate on, beyond which there is concern of the **memory resource** and **I/O constraints**, resulting in generation of ‘core’ file due to memory address violation error. Needless to mention, the varying sizes of different chromosomes would be causing improper work-load distribution and significant CPU wait time in case synchronization of output file generation and resubmission of left over job is desired. Hence, there is an acute need for proper synchronized distributed processing, optimizing the resources of a cluster such as the memory.

Now, let us consider two sequences of sizes N1 and N2 respectively. Time taken would be O (N1 * N2). For example, if N1=N2=100, time = O (100 * 100) = O (10,000)

Now let us fragment the two sequences into smaller sequences N1’=N1’’=N2’=N2’’=50 as in figure 6.

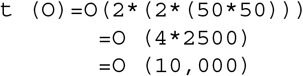

**Figure 5.**
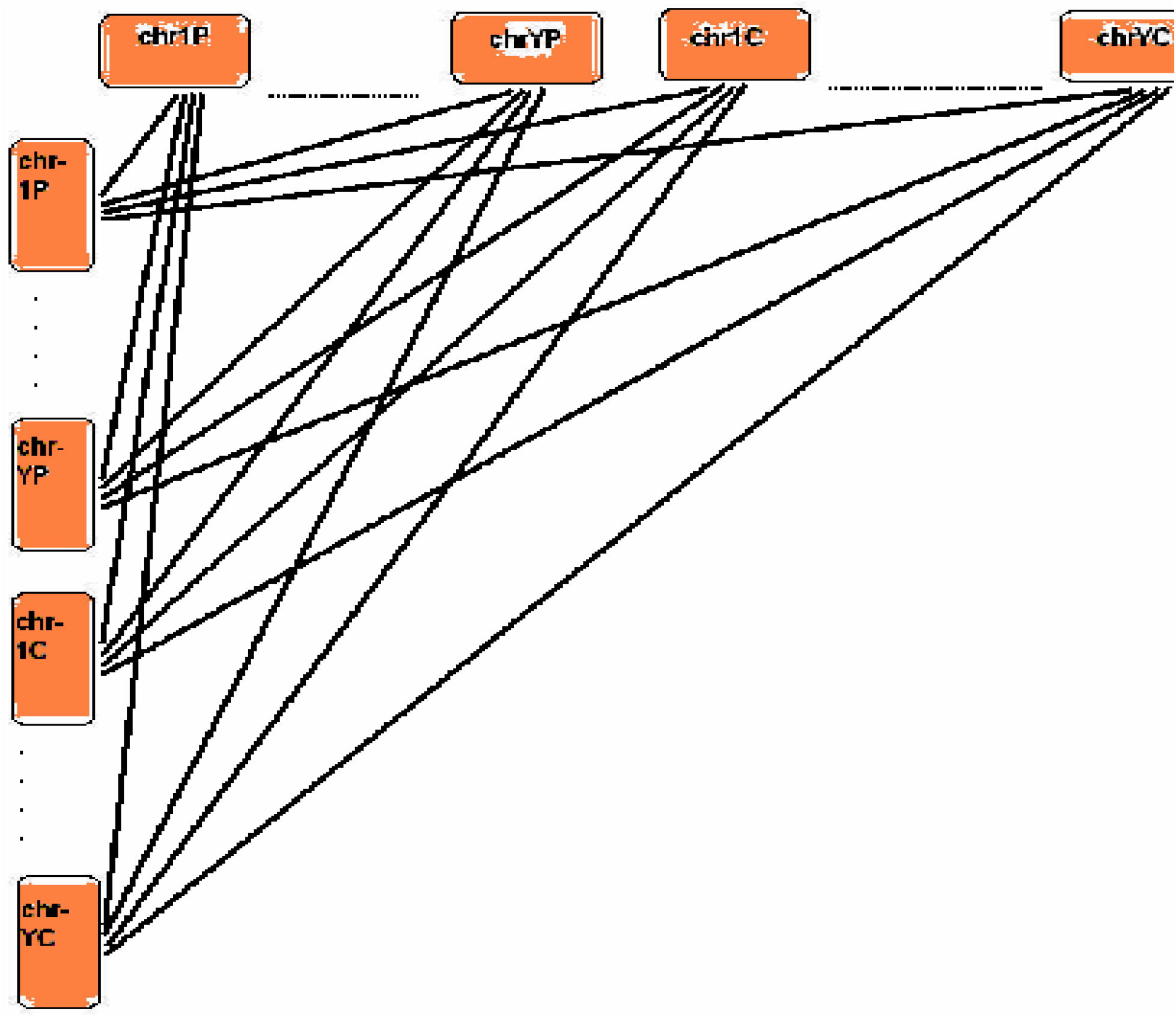

**Figure 6.**
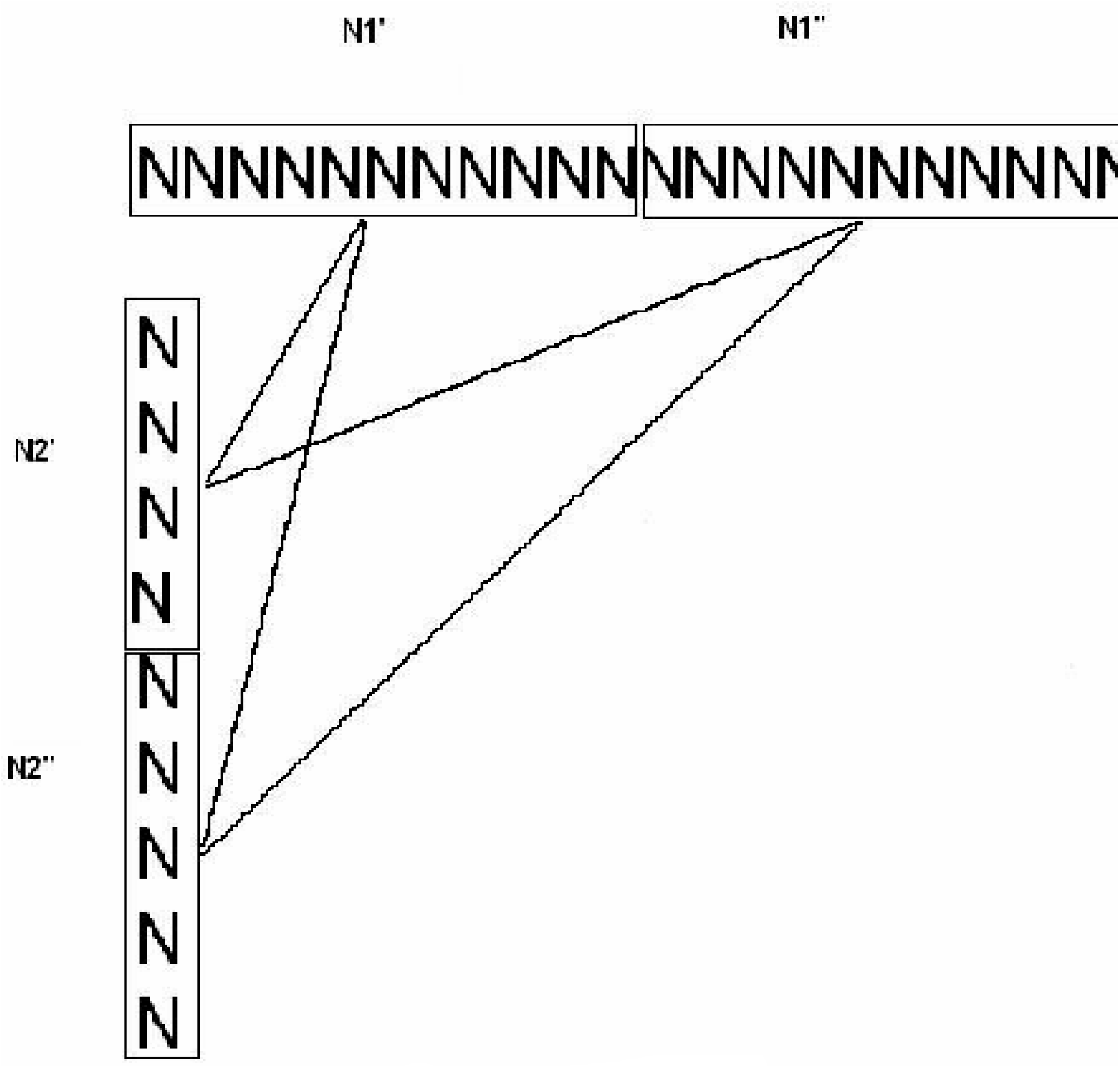

However, if we can distribute these ***independent comparisons*** into four Processors, then *ideally*,

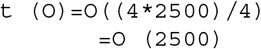

However, the splitting region where the partitioning was done can suffer, in the sense that we may miss matches at the site. The question is, ‘Do we really miss out the matches that we are interested in?’. The answer is ‘No’! The strong heuristic tools for matches such as blast, is so good in finding out a match that it can detect a good match of even as low bases as around 24 bases. Below is a graphical representation of the matches that can happen at the partitioning junction so that we can reverse engineer to stitch back the results generated as in figure 7.

**Figure 7.**
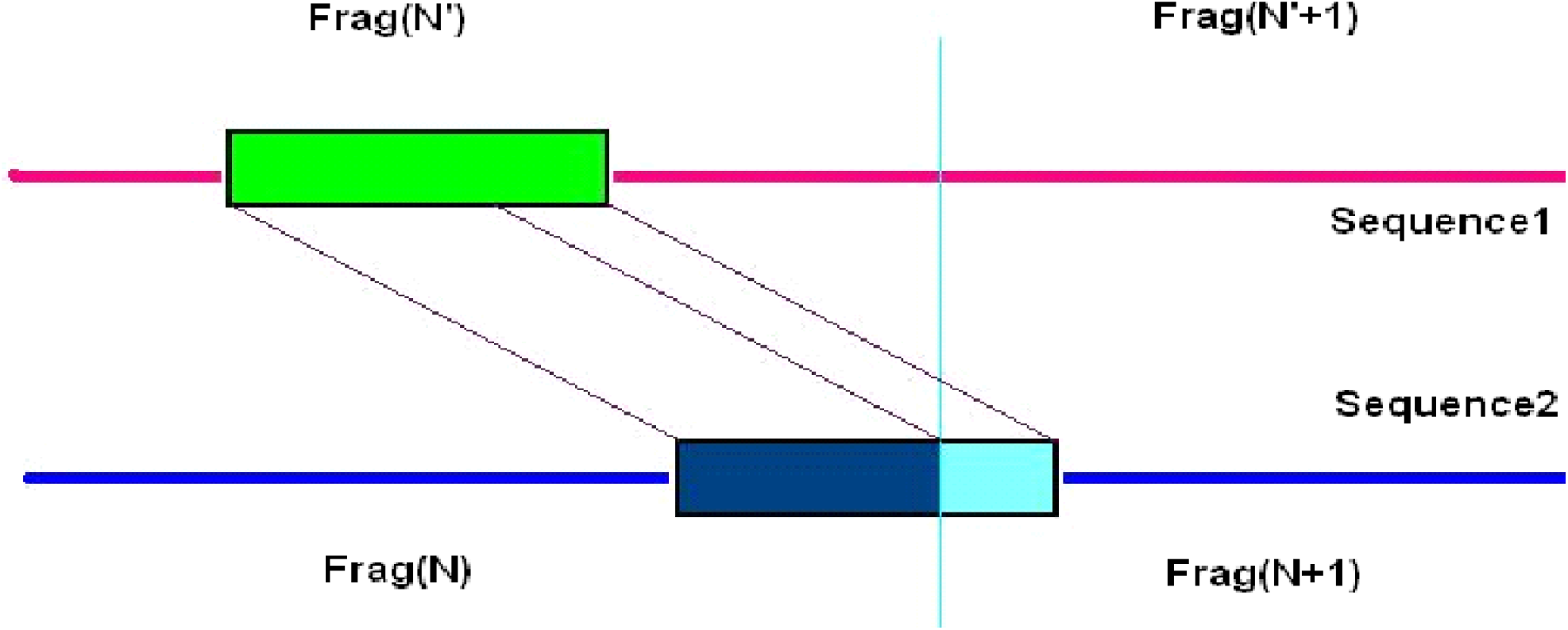

Of course, we would be missing matches of small number of nucleotides less than around 24 bases at the junction and one is interested in getting those *rare match events*, one can definitely pick up small nucleotides around the partition and compare it with all fragments for possible matches. The event would be *rare* since we would be using much higher fragment size relatively in the order of millions of bases. We choose to save time rather than identifying the rare events for the time being. Future version of the software might consider taking care of the *rare event* in case such a necessity comes up.

Combining the fragmentation concept with the genome alignment strategy, we would result in the following comparison algorithm as in figure 8.

**Figure 8.**
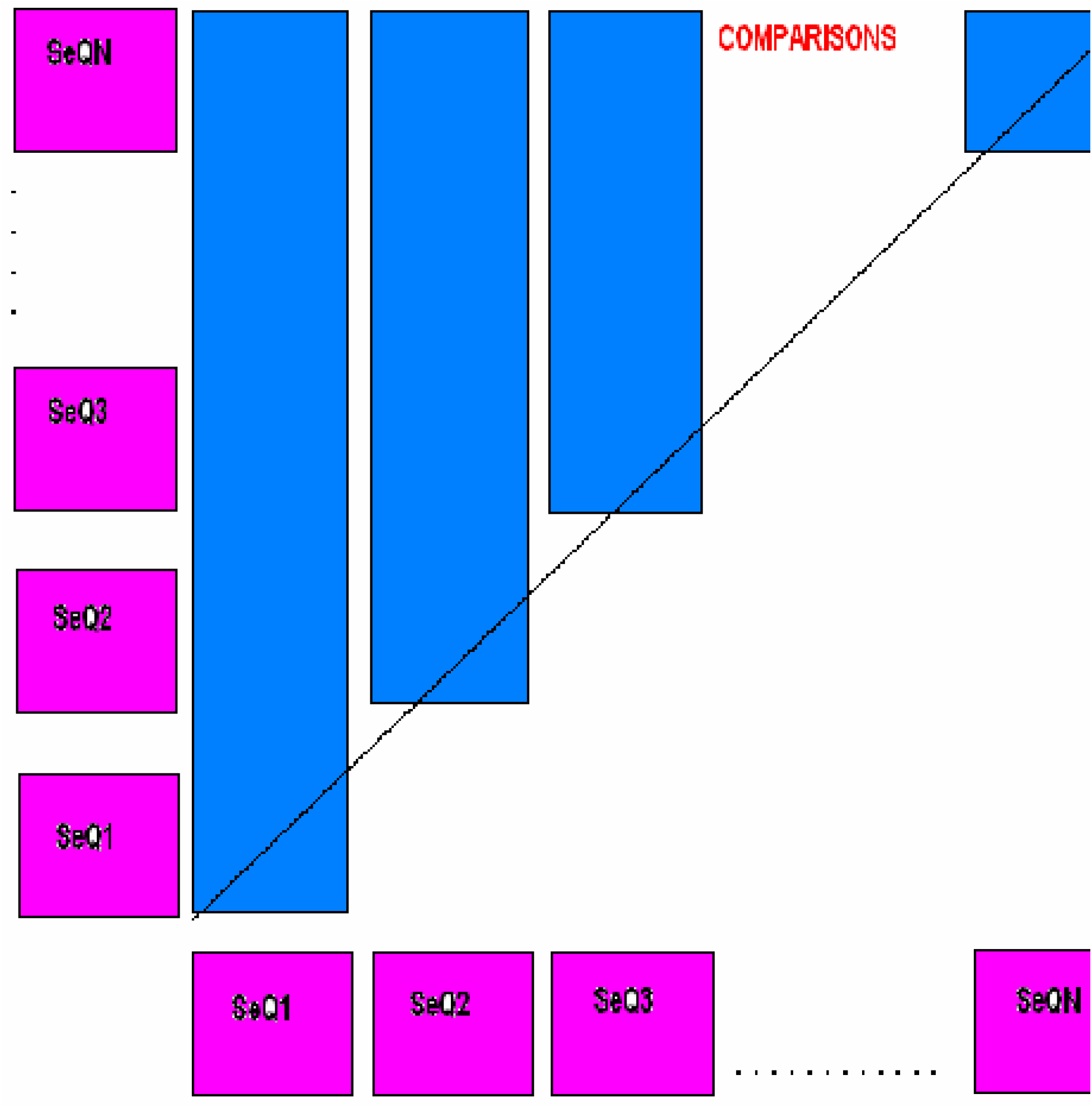

Where, SeQ1 to SeQN are the fragments of all the chromosomes of the two genome sequences in the order as stated in the genome alignment strategy. It is to be noted that by efficient programming we can quickly fragment the whole genome in the time order of few minutes.

Total comparisons for 1 fragment = N + (N − 1) + … + 3 + 2 + 1 = Summation N

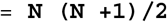

### Parallel Processing

Our goal should be to extract the information in a robust way and as rapidly as possible. The robustness can be introduced by automating the comparisons by means of parallel processing. We did the parallelization by means of message passing interface or **MPI**, though one can make use of other possible parallelizing tool for different architecture such as the symmetric multi-processor, or can even opt for serial job submission. MPI is a widely and uniformly accepted application programming interface, and an MPI script can be executed on any cluster for distributed job. The following scheme below falls under the category of *‘embarrassingly parallel’* programming. Intensive inter-processor communication is not required for the genome fragmentation and alignment strategy that we discussed above. One could also submit the jobs in a serial fashion. We wanted to make general purpose software to be operational on any cluster since the mode of job submission of different clusters is different, and thus general purpose serial job submission software is not possible because of lack of uniformity. Further, on most clusters an MPI script job has a higher priority over a serial job submission, and we wanted to take advantage of this fact, rather than requesting the cluster administration to change the priority settings.

Since the fragments are of similar size, the job execution time on each node would be comparable such that there would not be a significant idle time of any processor at the barrier as shown in figure 9. GMSECT, works best while comparing genomes of two individuals from the same organism such that the data quality is same thereby further facilitating reduced idle time of any processor at the barrier. Of course, ‘GMSECT’ can be used for comparing distant species as well, though in that case the idle CPU time might increase a bit. Future version of the software might consider taking care of this aspect as well in case such a necessity comes up by introducing separate barriers at each node. It is to be noted that introducing barrier at each node can greatly increase the latency due to increased calls from the master to the slaves, causing a drop-down in the time performance in case comparing closely related species.

**Figure 9.**
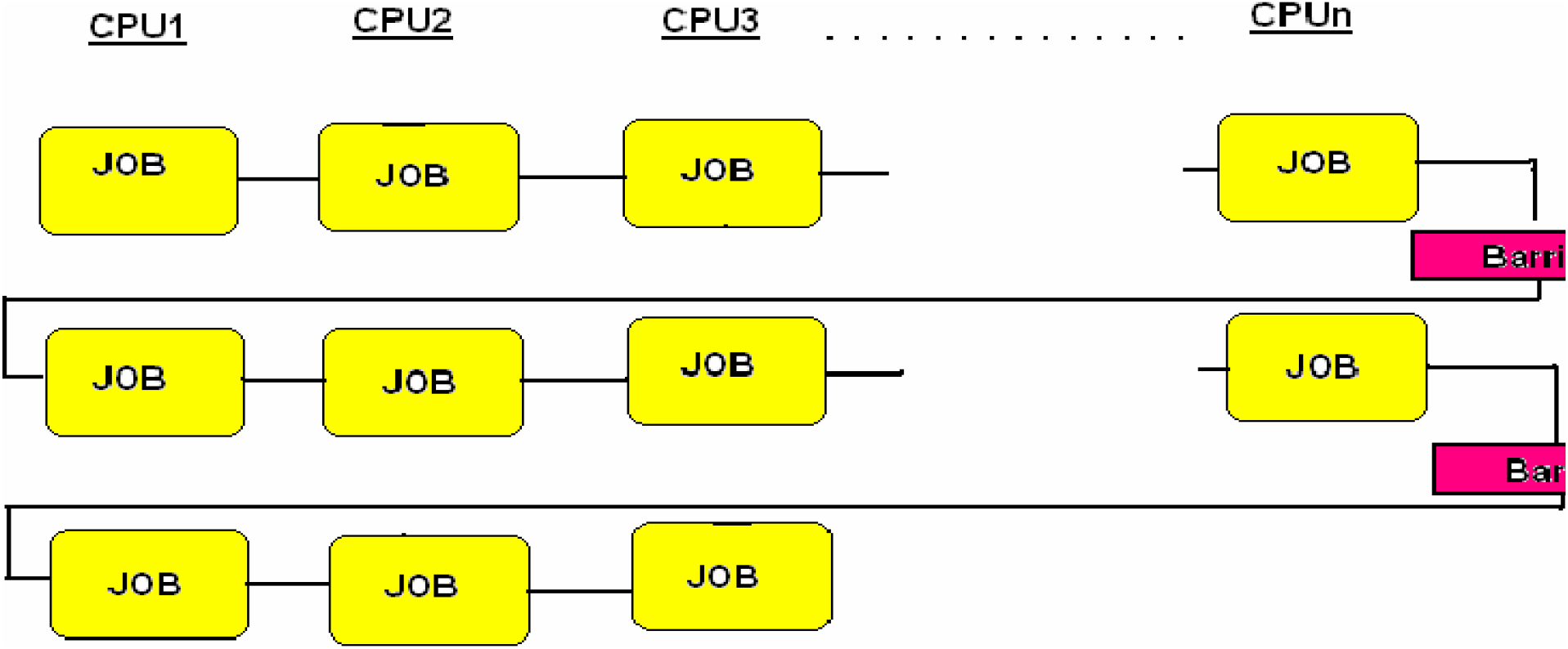

Average Work Load on each Processor for 1 fragment = **N (N+1)/2n**, where ‘n’ is the number of processors allotted.

### Optimization

With the vision of the personalized genome sequencing project we realize the amount of data that would be generated and the huge amount of information that can be extracted from each genome, requiring us tremendous and efficient computational requirements. Hence, for mass comparison of 100s of individual’s genome, or even otherwise, we would prefer to maximally utilize the resources of a cluster in order to minimize the execution time. The secondary concern we had was to minimize the partitioning required so that there is minimum requirement of stitching back the matches, and to make the small base matches at the partitioning junction as a rare event, contributing to negligible information that would be lost in comparison to huge information that would be obtained.

In figure 10 is the profile of fragmentation size versus comparison time using GMSECT operated on chromosome 21 of Celera’s Human Genome compilation in a self comparison with 15 processors each with 2GB memory and 2.2 GHz using blast.

**Figure 10.**
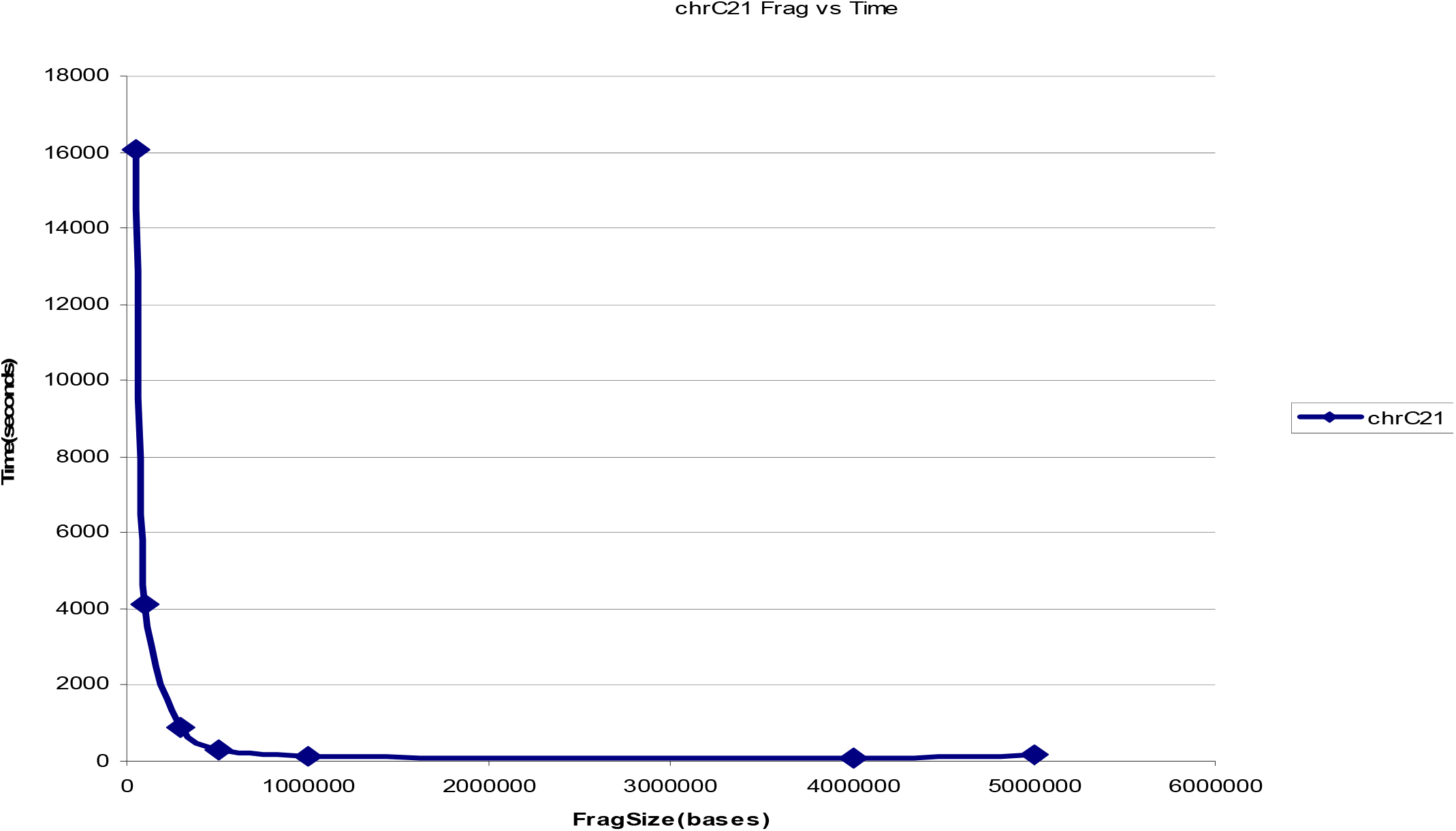

The profile shows hyperbolic nature resemblance till a fragmentation size of 4 million bases, beyond which the curve shoots up. In other words, the Minima is obtained at 4 million bases under the given conditions. Further, creating fragment size of more than 5 million bases resulted in generation of ‘core’ files.

A similar hyperbolic resemblance profile was generated when GMSECT was operated on a distant species to human such as chromosome 2 of *Arabidopsis thaliana* under the same conditions using the blast tool choice as shown in figure 11.

**Figure 11.**
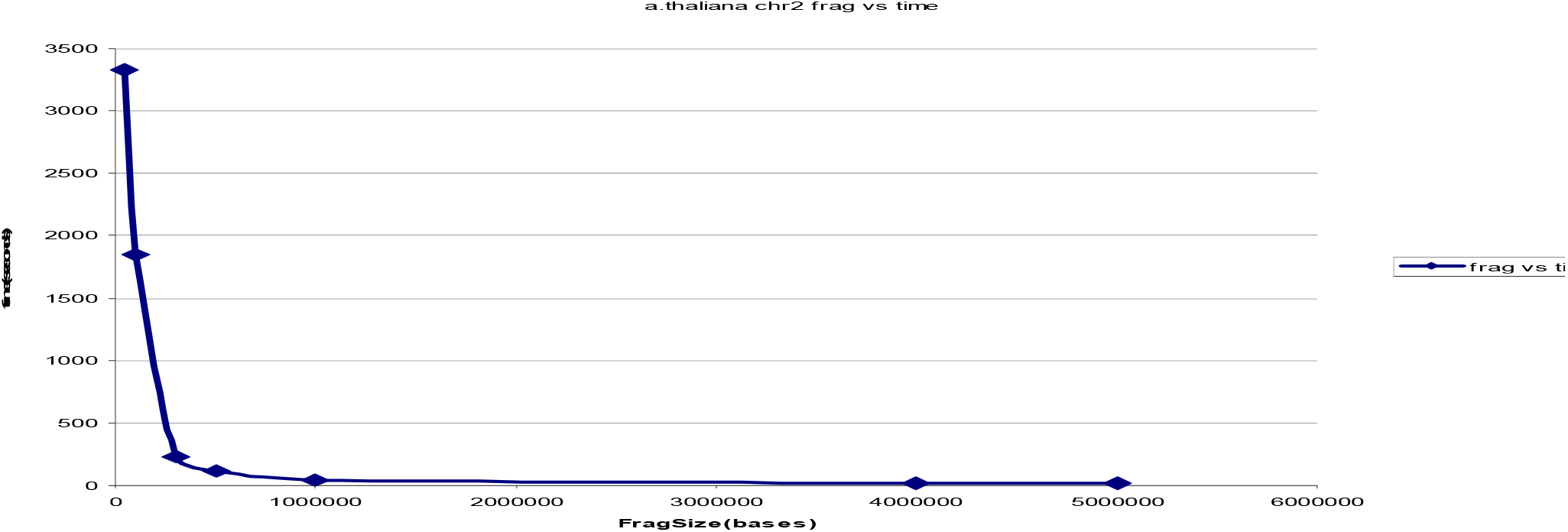

The Minima was found again at around 4 million bases. The slight shift of minima could be attributed to the change in data quality. Here again, ‘core’ files were generated when fragment size of 5 million or more was used due to memory address violation error.

Since the Minima is around 4 million bases, even for distant species, we can safely consider this value to be a ‘general’ **optimal fragmentation size** on HPF cluster nodes each having processor of the type mentioned earlier. Reducing the fragmentation size to smaller size would not only increase the computational time, but also increase the approximation of stitching back the matches at the junction.

### Performance and Evaluation

It is to be noted that with the increase in the sequence size, the number of comparisons increases by a power of two, because of the very nature of ‘Summation N’ formula i.e. N (N+1)/2.

If x = N (number of fragments) and y = number of comparisons, Then y = x(x+1)/2

This equation is of the form **X^2 = 4aY**, with X & Y considered are variable with frame-shift of x & y respectively. Thus, the concern is how the GMSECT performs with regard to increase in comparison jobs. In figure 12 is the number of comparisons versus time graph generated using increased units of chromosome 21 units of Celera’s Human Genome compilation using the blast tool choice with standard output format.

**Figure 12.**
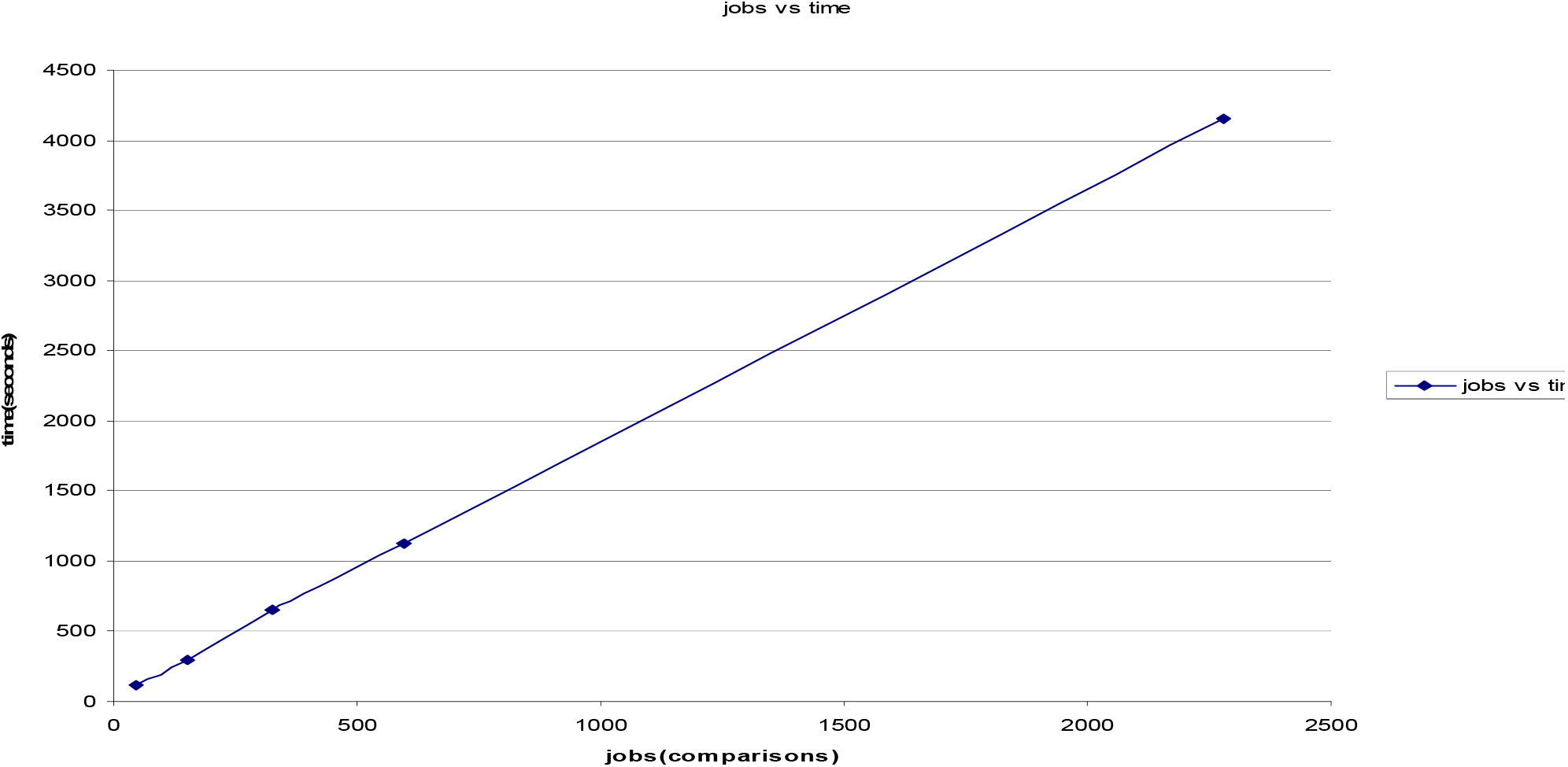

The above **superlinear** behavior is a positive indication of the trust on the performance.

In figure 13 are the processors versus time performance curve which indicates **high scalable performance** using GMSECT on chromosome 21 versus itself of the Celera’s Human Genome compilation using the blast choice option having standard output format.

**Figure 13.**
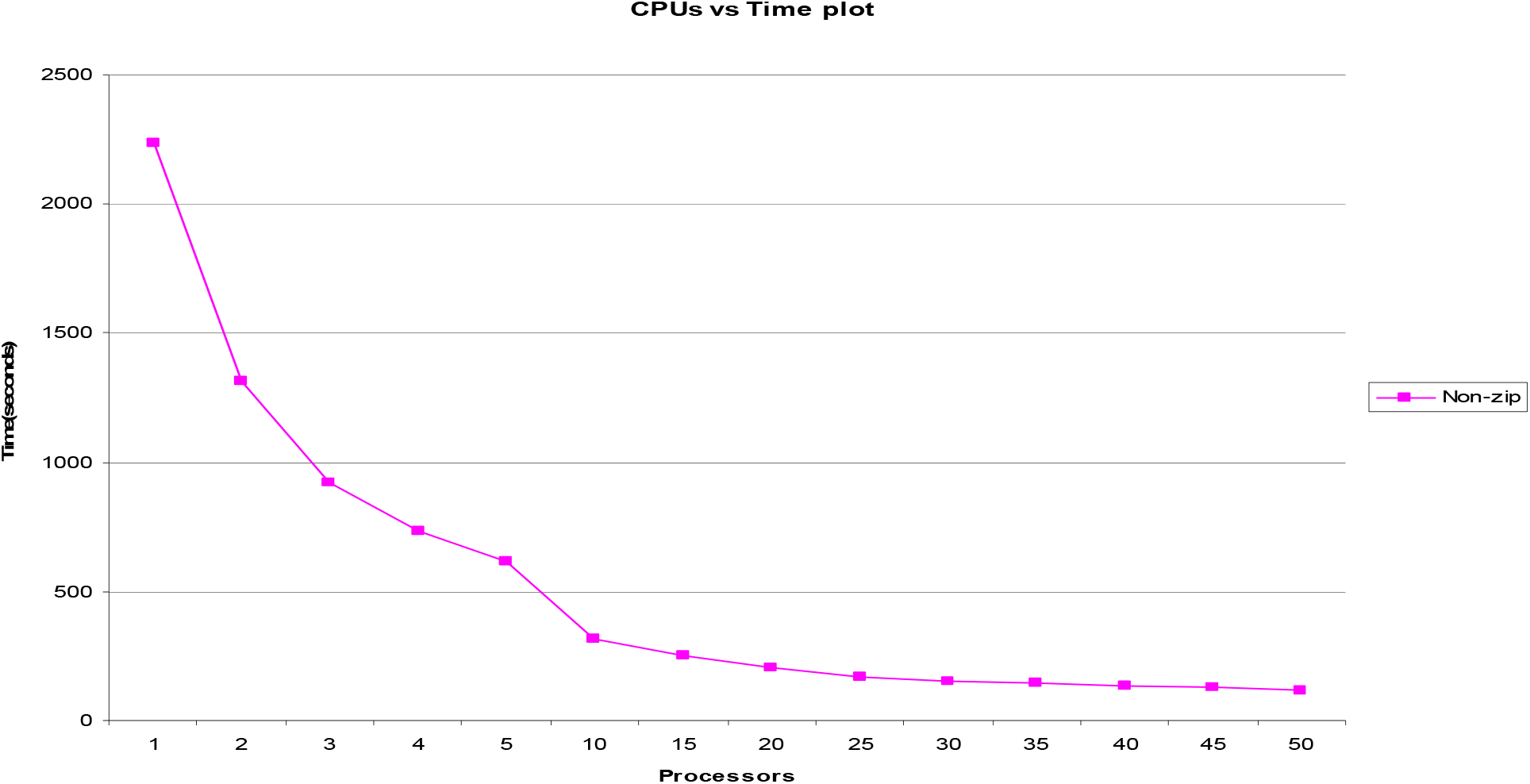

### Sample Execution

We compared the chromosome 21 of Celera’s human genome compilation with all of its 24 chromosomes and with all the 24 chromosomes of the human build 35 reference sequence using 110 processors of HPF each with 2GB memory and 2.2 GHz capacity, using the pairwise blast comparison heuristic choice with all alignments having statistical relevance of expectation value (e-value) less than 1 and the tabular output format. The comparison was completed in just 2 hours and 10 minutes! Using the above information it is estimated that comparing two unique human genomes would take around 9.4 days comprising of two self comparisons and a non-self. GMSECT can even be applied to microbial genome such as the *Escherichia coli* or algae *Botryococcus braunii* or yeast *Saccharomyces cerevisiae* to quickly do the comparisons, and thus finds its application to the pharmaceuticals and microbial product based firms for the research and development.

## Future Work & Supplementary Materials

There is scope of developing a new version of GMSECT that could take care of comparing contigs rather than comparing sequences, since not all contigs are mapped to the chromosomes. Further, the code can even be developed to take care of matches of around 24 bases or less at the splitting junction which we have not bothered at this stage due to the even being a rare event due to extremely large fragment sizes in the order of millions of bases. Supplementary materials can be downloaded from https://sites.google.com/a/iitdalumni.com/abi/educational-papers.

